# Optimal targeted therapy for multiple cancers based on contrastive Notch signaling networks

**DOI:** 10.1101/2024.06.26.600739

**Authors:** Tamaki Wakamoto, Sungrim Seirin-Lee

## Abstract

Over decades, cancer understanding has advanced significantly at molecular and cellular levels, leading to various therapies based on intra-/inter-cellular networks. Despite this, cancer still remains a leading cause of death globally. The primary driver of cancer mortality is metastasis, responsible for about 90% of cancer deaths, due to unclear pathophysiological mechanisms that complicate treatment development. The Notch signaling pathway, a crucial intercellular network in many cancers, has been extensively studied and therapies targeting the Notch pathway also have been well-studied based on inhibiting various stages of Notch activation. On the other hand, Notch signaling’s role varies between cancers; for instance, in non-small cell lung cancer, Notch1 and Notch2 have opposing effects compared to their roles in embryonal brain tumors. In this study, we assumed a scenario of multiple cancers with contrasting Notch signaling pathways and explored optimal targeted therapies for reducing cancer cells by developing two mathematical models with contrasting Notch signaling pathways. The proposed therapies were compared with existing ones, and strategies were investigated to reduce cancer cell numbers for different stage of cancer. We found that that multiple cancers with contrasting Notch networks can be controlled by a common targeted signal network. Combination therapy enhancing Notch production may be most effective in early-stage cancer, while cleavage therapies may be more effective in late-stage cancer. Our study also suggests that optimal treatment should consider the cancer stage, with careful selection and ordering of medication therapies.

## 1 Introduction

Over decades, the understanding of cancer has been dramatically progressed in molecular and cellular levels and the treatments based on intra-/inter-cellular networks have been widely developed (Hassanpour and Dehghani 2017; Bidram et al. 2019). Nonetheless, cancer is still a life-threatening disease and becomes a major cause of death worldwide (Sung et al. 2021). According to the recent statical study (Sung et al. 2021), 19.3 million new cancer cases and 10 million deaths resulted from cancer have been reported in the world. Notably, the major cause of cancer mortality is not primary cancers but cancer metastasis which account for about 90 % of cancer deaths (Guan 2015; Lambert et al. 2017). The reason why cancer metastasis is the principal cause of cancer lethality is due to the fact that the pathophysiological mechanism of metastasis *in vivo* remains unclear, making it difficult to develop appropriate treatments and medications (Suhail et al. 2019; Fares et al. 2020).

The Notch signaling pathway is an important intercellular network common to several cancers (Yuan et al. 2015; Venkatesh et al. 2018; Zhou et al. 2022), which has been also well-studied as a cell fate-determining network during development process (Hori et al. 2013). The Notch signaling pathway starts with the combination of the transmembrane receptor Notch and its ligand Delta on the cell membrane (Fig. 1A). The binding of Notch in one cell and Delta in adjacent cells leads to the release of the Notch intracellular domain (NICD) into the cytosol. Through signaling pathways, it binds to transcription factors, resulting in the activation of Notch production and the inhibition of Delta production via HES-1 gene expression (Boareto et al. 2015).

**Figure 1:**
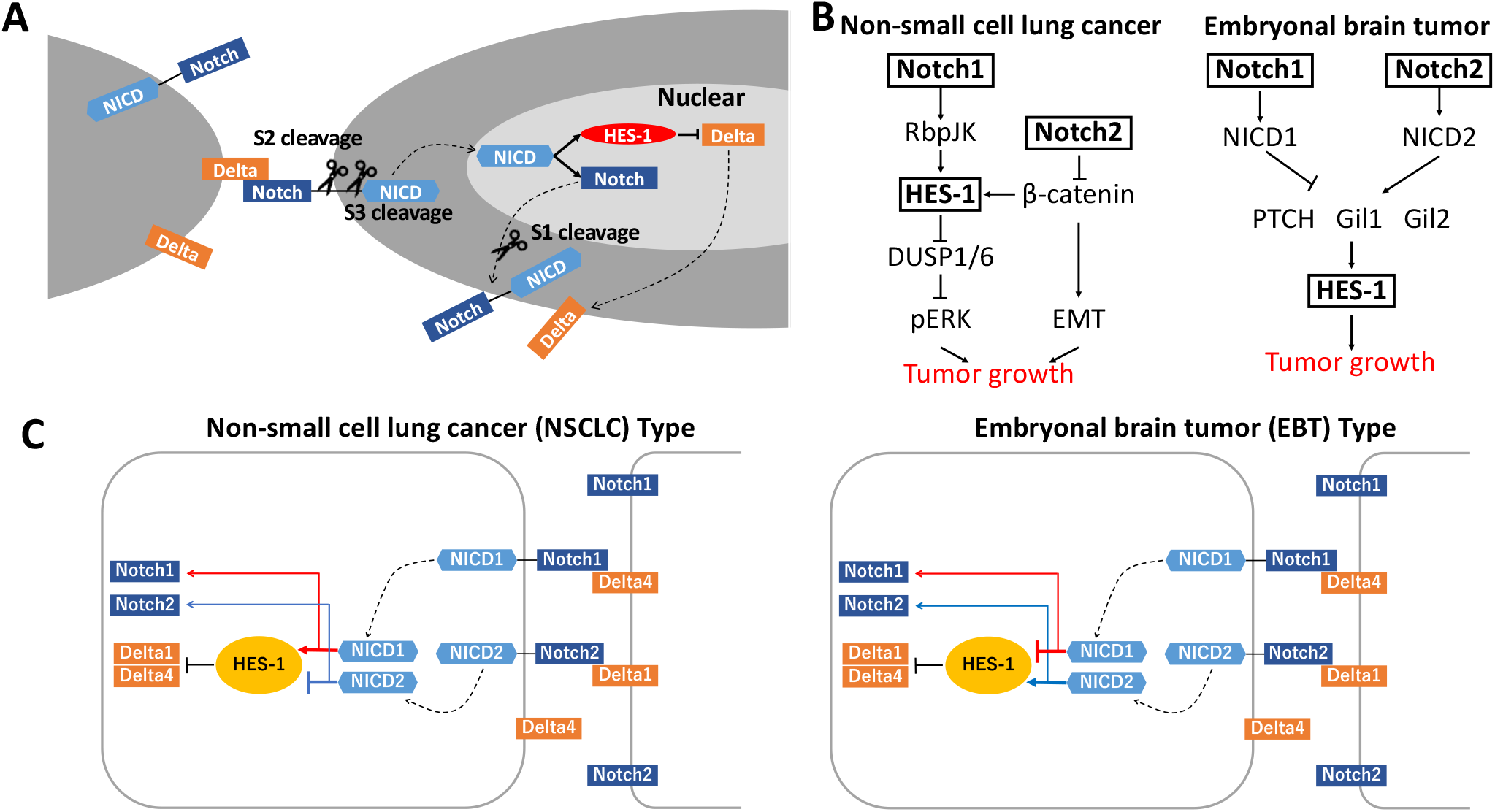
Network of Notch signaling. (A) Diagram of Notch signaling network. Dot arrows represent the translocation of the biochemicals. (B) Diagrams of the gene network in non-small cell lung cancer and embryonal brain tumor. (C) Diagrams of Notch signaling network in non-small cell lung cancer and embryonal brain tumor.

Various cancer therapies such as chemotherapy, hormone therapy, immunotherapy, radiation therapy, surgery, and targeted therapy, are being implemented in clinical settings depending on cancer types (Bidram et al. 2019). In particular, targeted therapy is one of precision medicine types that targets proteins controlling the networks of cancer cell growth, proliferation, and spread (Min and Lee 2022; Zhou and Li 2022). The conventional targeted therapy designs DNA changes or protein targets using a specific inhibition signal transduction pathways. Various therapies targeting Notch signaling have been well-studied and tested in preclinical and clinical settings. The therapies are based on the methods that either inhibit a cytoplasmic path producing the Notch receptors on cell membrane (S1 cleavage in Fig. 1A) or inhibit Delta-Notch binding (S2 cleavage or S3 cleavage in Fig. 1A) (Yuan et al. 2015; Venkatesh et al. 2018; Zhou et al. 2022). S1 cleavage is involved in the process that the precursors of Notch to become mature Notch and translocate to cell membrane (Zhou et al. 2022). S2 and S3 cleavages are both required to NICD release. S2 cleavage occurs outside of cell when Delta binds to Notch but S3 cleavage detaches NICD from the Notch receptor by *γ*-secretase in cytosol (Mumm et al. 2000; Golde et al. 2013; Zhou et al. 2022).

Non-small cell lung cancer (NSCLC) is a critical primary tumor to occur brain tumor metastases (Baumgart et al. 2015; Nishino et al. 2019). About 40 to 50 % of cases with lung cancer have brain metastases and approximately, 10-20% of NSCLC patients present brain metastases (Nishino et al. 2019). The conventional pathway of cancer growth by Notch signaling is based on activating oncogene HES-1 regulated by notch receptors, Notch1 and Notch2 (Fan et al. 2004). For example, in NSCLC, Notch1 activates HES-1 via RbpJK while Notch2 inhibits HES-1 via *β*-catenin (Baumgart et al. 2015) (Fig. 1B, left panel). In contrast, embryonal brain tumor (EBT) has an opposite pathway. Notch1 intracellular domain (NICD1) inhibits HES-1 while Notch2 intracellular domain (NICD2) activates HES-1 via Hedgehog pathway, PTCH, Gil1, and Gil2 (Fan et al. 2004) (Fig. 1B, right panel). That is, Notch1 and Notch2 have contrasting roles between NSCLS and EBT (Fig. 1C). This implies that if we enhance Notch1 to inhibit HES-1 gene and suppress EBT growth, it may result in up-regulation of HES-1 gene in NSCLC. Conversely, if we enhance Notch2 to inhibit HES-1 and suppress NSCLC growth, EBT will be worsen. In other words, if metastasis occurs in multiple cancers with a constitutive Notch network, targeted therapy for the Notch network should be very carefully considered.

Therefore, in this study, we focus on developing targeted therapies to suppress multiple cancers with contrasting Notch signaling networks. To this end, we developed two mathematical models representing these contrasting Notch pathways and explored the existence of a common signaling pathway that could simultaneously suppress both cancers. Using results from network and sensitivity analyses, we suggest new targeted therapies and compare their effectiveness in reducing cancer cell numbers with existing therapies. Additionally, we investigate how the order of therapy combinations is important in increasing the effectiveness of therapies and what is the optimal strategies to decrease both cancer cell numbers and expansion speed in Stage 4 cancer progression. Our results suggest that multiple cancers with contrasting Notch networks can be controlled by a common targeted signal network. Combination therapy enhancing Notch production may be most effective in early-stage cancer, while cleavage therapies may be more effective in late-stage cancer. Optimal treatment should consider the cancer stage, with careful selection and ordering of medication therapies.

## 2 Method

### 2.1 Basal Models of Notch signaling: NSCLS type and EBT type models

We develop two Notch signaling models with contrastive roles of Notch1 and Notch2. The network pathways of Notch signaling are described in one/two dimensional layer/surface of cell membrane and two/three dimensional bulk space of cell cytosol as shown in Fig. 1C. Since we focus on the kinetic dynamics of molecular networks, we neglect the details of cell geometry and choose the model reduction approach to cell geometry-free type of Notch signaling suggested in Seirin-Lee (2016). Further assumptions for models are given as the following.

- Cells are arranged on a grid {(*i, j*) | 1 ≤ *i* ≤ *m*, 1 ≤ *j* ≤ *n*} and Notch receptor on one cell binds to Delta ligands from four adjacent cells (Fig. 2A).
- The signal by Notch-Delta binding is transferred as the average level from adjacent cells (Collier et al. 1996).
- We simplify Notch signaling to only include the Notch1 and Notch2 pathways, focusing solely on canonical Notch signaling pathways (Kopan and Ilagan 2009).
- In NSCLC type model, the gene expression of HES-1 is decreased by the level of Notch2 intracellular domain (NICD2) and increased by the level of Notch1 intracellular domain (NICD1) (Baumgart et al. 2015) (Fig. 1B).
- In EBT type model, the gene expression of HES-1 is increased by the level of NICD2 and decreased by the level of NICD1 (Fan et al. 2004) (Fig. 1B).
- The same signaling proteins have the same decay, transcript, and transport rates.

**Figure 2:**
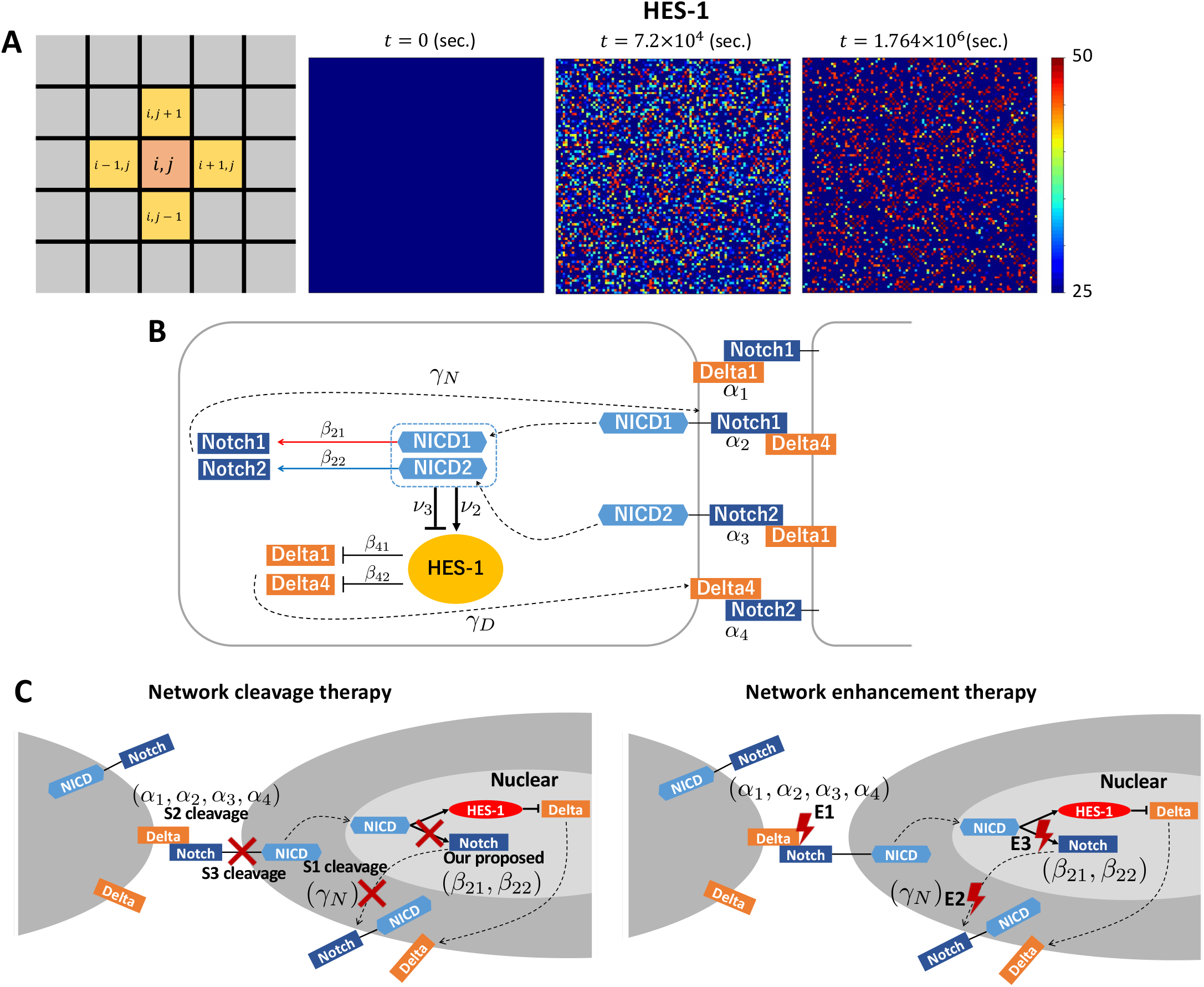
Simulation method and example. (A) Cell position and pattern of HES-1 expression. A cell (*i, j*) is surrounded by four cells ((*i, j* + 1), (*i, j* − 1), (*i* + 1, *j*), and (*i* − 1, *j*)) and interacts with these cells. (B) Notch signaling and corresponding parameter variables in Eq. (1). (C) Diagram of the Notch signaling network with therapies. Network cleavage therapy is marked by red cross. Network enhancement therapy is marked by red lightning.

For each (*i, j*)-th cell (1 ≤ *i* ≤ *m* and 1 ≤ *j* ≤ *n*), let us denote the concentrations of Notch1 (N1) and Notch2 (N2) in cell membrane and cytosol by 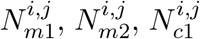, and 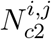, respectively, the concentrations of Delta1 (D1) and Delta4 (D4) in cell membrane and cytosol by 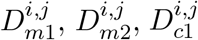, and 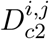, respectively, and the concentrations of NICD1, NICD2, and HES-1 by 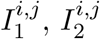, and *H*^*i,j*^ respectively. Then, the model equations on (*i, j*)-th cell are given to

#### Notch in membrane

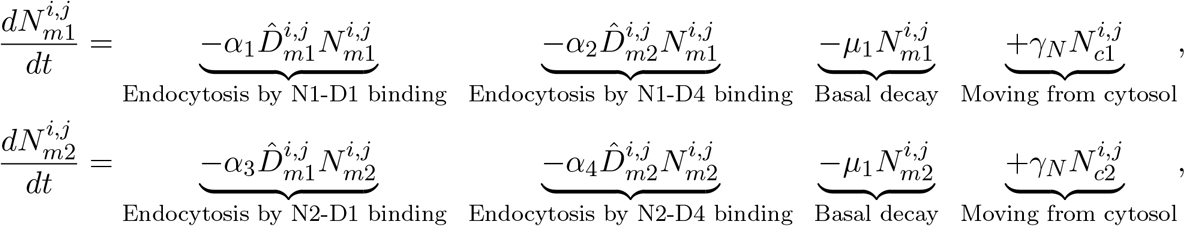

#### Delta in membrane

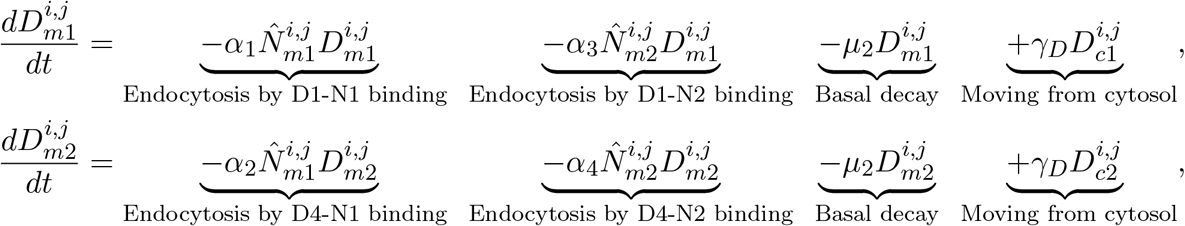

#### NICD in cytosol

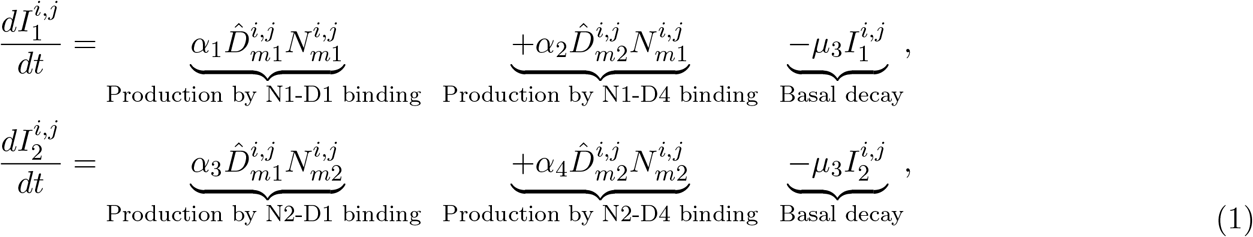

#### HES-1 in nucleus

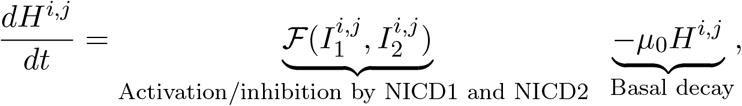

#### Notch in cytosol

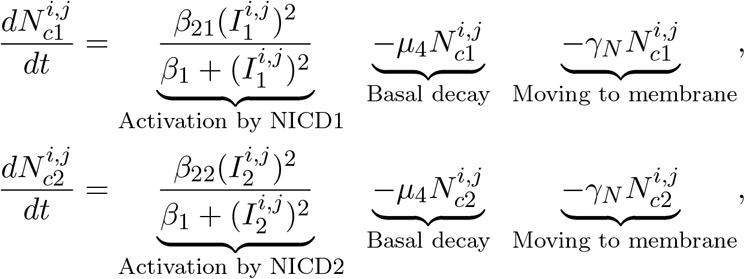

#### Delta in cytosol

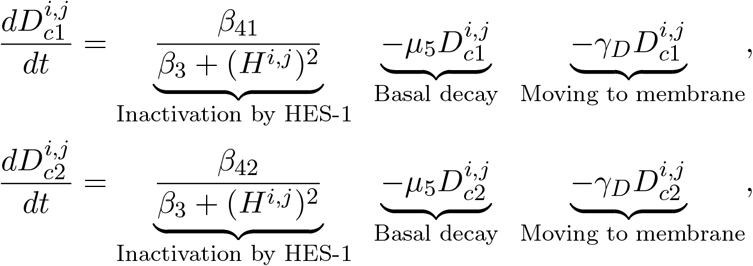

where *α*_1_, *α*_2_, *α*_3_, and *α*_4_ are the binding rate of Notch1-Delta1, Notch1-Delta4, Notch2-Delta1, and Notch2-Delta4, respectively (Fig. 2B). *µ*_0_, *µ*_1_, *µ*_2_, *µ*_3_, *µ*_4_, and *µ*_5_ are basal decays. *γ*_*N*_ and *γ*_*D*_ are translocation rates of Notch and Delta, respectively (Fig. 2B). 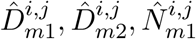, and 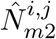 are the average concentrations of Delta1, Delta4, Notch1, and Notch2 in the cell membrane adjacent to (*i, j*)-th cell, respectively, and denoted as follows:

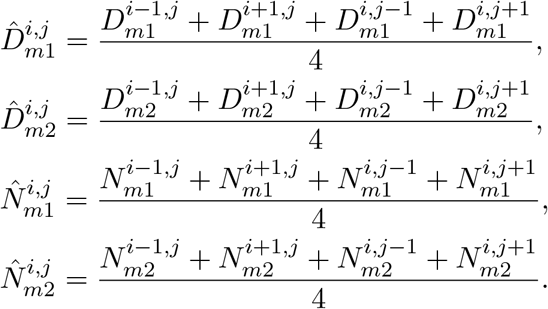

We model 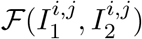 as a function of NICD1 and NICD2 in NSCLS type case and EBT type case, respectively.

#### NSCLS type model

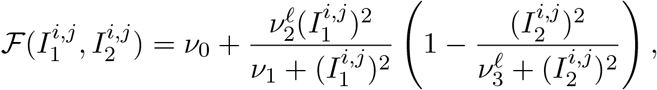

#### EBT type model

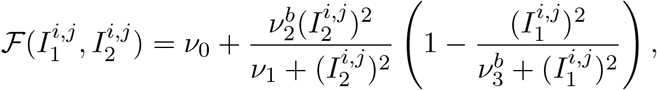

where *ν*_0_ is the basal expression rate and 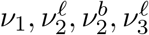 and 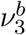 are the positive constants regulating activation and inhibition of HES-1 expression.

### 2.2 *in silico* targeted therapy model I: Network-cleaved therapy

In this study, we will test two targeted therapies suggested in Pine (2018), Pagliaro et al. (2021), and Lu et al. (2019), and a new targeted therapy. (i) S1 cleavage (denoted by the parameter 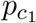) which inhibits a cytoplasmic path producing the Notch receptors on cell membrane. (ii) S2 & S3 cleavages (denoted by the parameter 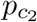) which inhibit Notch–Delta binding in cell membrane. These two existent targeted therapies, (i) and (ii), are shown in Fig. 2C. Note that we model S2 and S3 cleavages by a single parameter because both S2 cleavage and S3 cleavage are required to block NICD release (Mumm et al. 2000). On the other hand, we will introduce a new targeted therapy based on our network analysis, which suppresses the production of Notch in cytosol by NICD. We denote its effect by the parameter 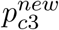. The *in silico* targeted therapy model is given to

#### Notch in membrane

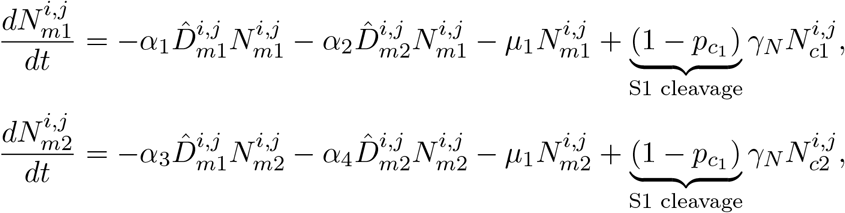

#### NICD in cytosol

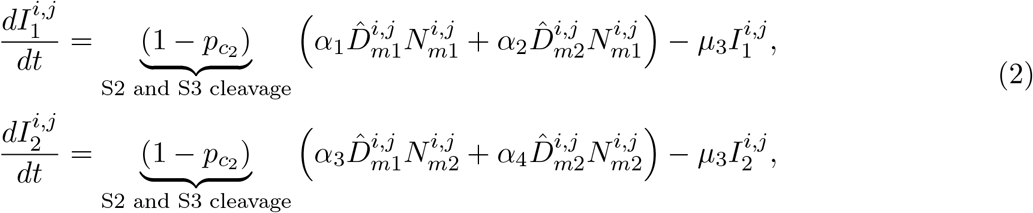

#### Notch in cytosol

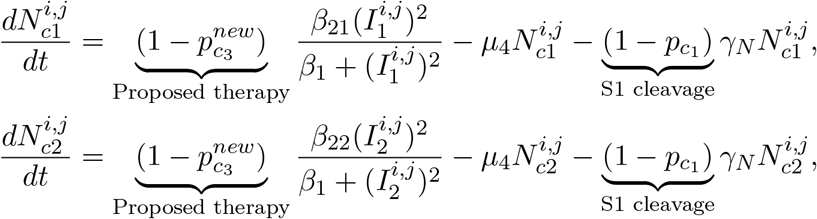

where 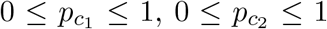, and 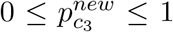 are the intensities of treatments targeting S1 cleavage, S2 and S3 cleavages, and Notch expression in cytosol, respectively. We chose these values as 10 random samples to consider the effective variance among patients.

### 2.3 *in silico* targeted therapy model II: Network-enhanced therapy

Based on our network analysis, we will additionally suggest a network-enhanced therapy. In this targeted therapy, we will enhance a targeted network as follows. (i) E1 enhancement (denoted by the parameter 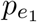) which enhances Notch translocation from cytosol to cell membrane. (ii) E2 enhancement (denoted by the parameter 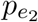) which enhances Delta-Notch binding in cell membrane. (iii) E3 enhancement (denoted by the parameter 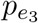) which enhances the production of Notch in cytosol by NICD.

#### Notch in membrane

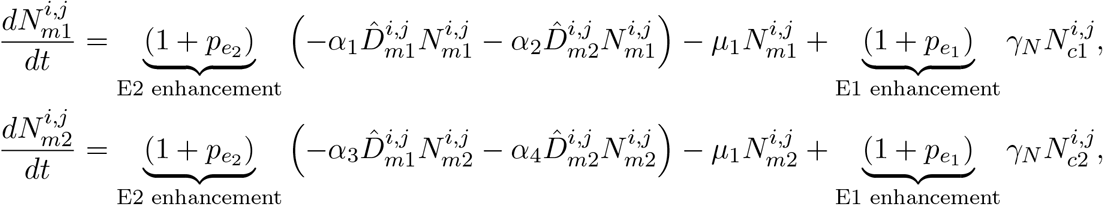

#### Delta in membrane

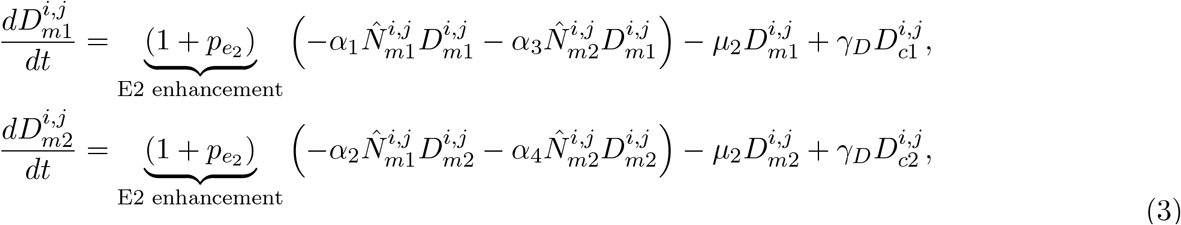

#### NICD in cytosol

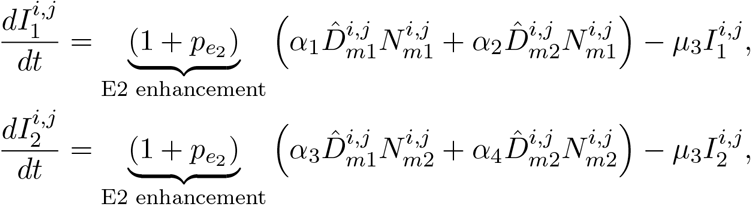

#### Notch in cytosol

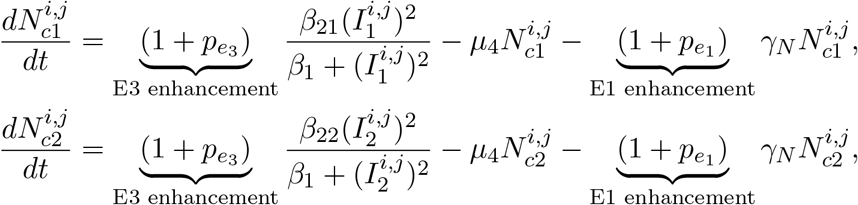

where 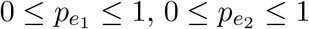, and 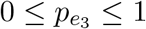 are ratio of treatment targeting Notch translocation to cell membrane, Delta-Notch binding, and Notch expression in cytosol, respectively. We chose these values as 10 random samples to consider the effective variance among patients.

### 2.4 Definition of cancer cells

In both NSCLC and EBT, cancer cells express oncogene HES-1 higher than healthy cells (Baumgart et al. 2015; Fan et al. 2004). Thus, we define a cancer cell by a cell expressing high level of representative oncogene (HES-1 in this study) such that

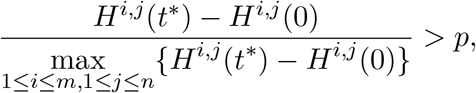

where *t*^*^ is the time when the solutions approached to a steady state. *p* is the indicator ratio of increased HES-1 expression compared to the initial level. We also define an index counting a total cancer cell number by

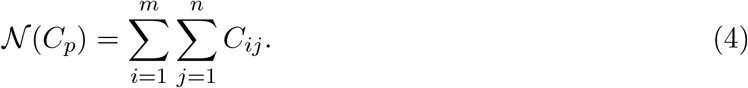

where

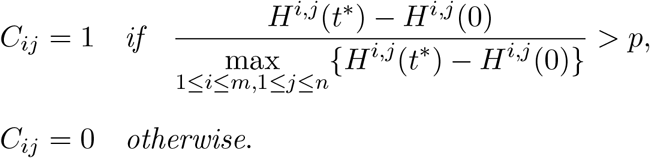

The index, *𝒩* (*C*_*p*_), gives us how many cancer cells with HES-1 expression over *p* × 100% from the initial level increased. We typically choose *p* =0.5 or 0.75.

### 2.5 Representative parameters, initial and boundary conditions, and sensitivity functions

We first assume that basal decay rates and translocation rates to the cell membrane are represented by the same parameters for similar types of proteins. On the other hand, we assume that the binding rate of Notch and Delta is different for each combination and is represented by different parameters. We also suppose that binding rates satisfy (Kakuda et al. 2020):

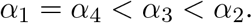

We chose a spatial scale and time scale as 6 cm and 3.6 × 10^3^ seconds that approximately corresponds to development time scale of lung cancer (Tubiana 1989). For the kinetic parameters, we chose representative values that generate a salt and pepper pattern. The details of parameter and values are summarized in Table 1.

**Table 1:**
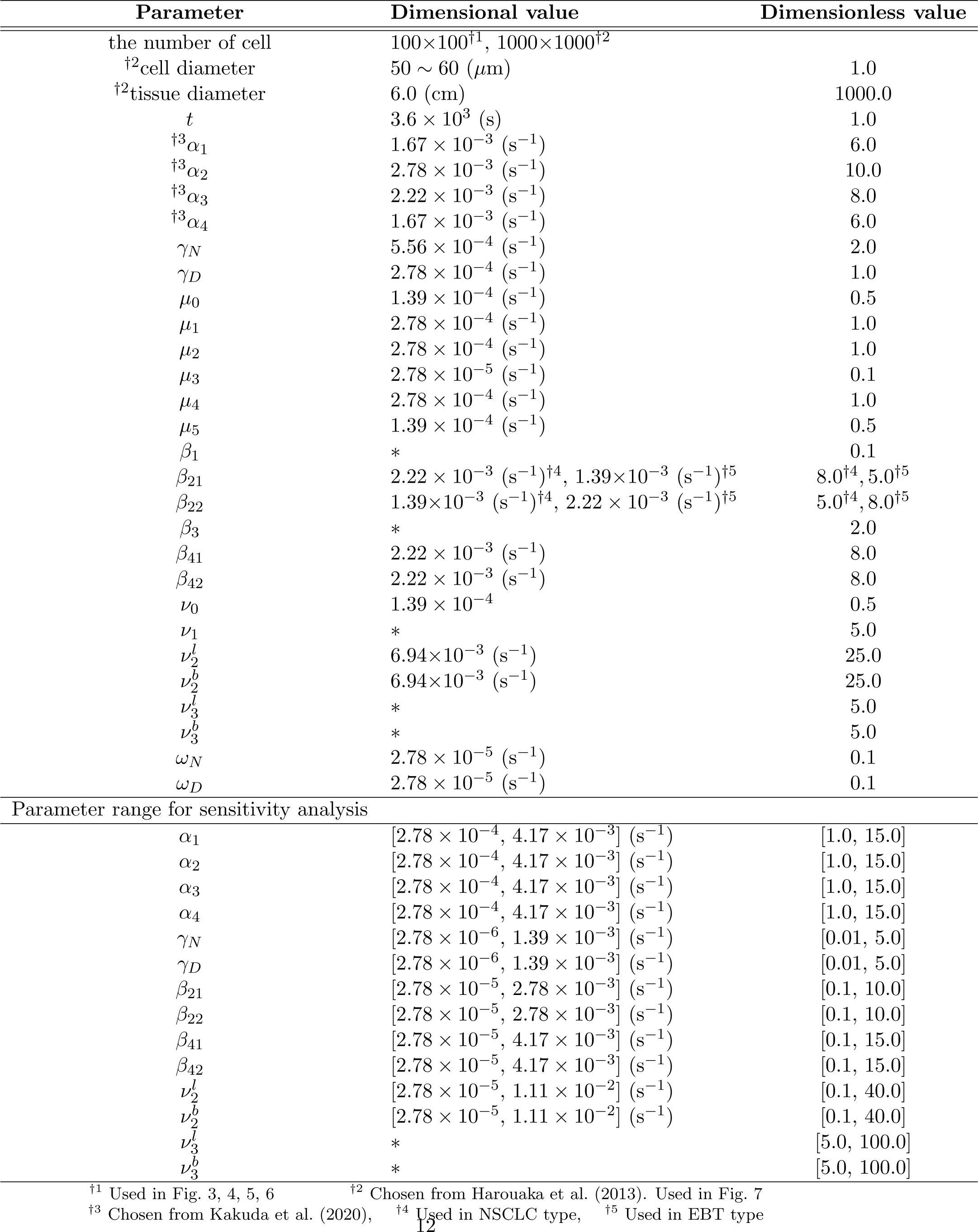
Representative parameter set.

We represent the cells as a regular array of identical square. 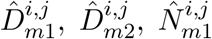, and 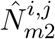 denote the mean over the four immediately neighboring cells for cell (*i, j*) (Fig. 2A). To consider the early stage and late stage of cancer dynamics, we assume the two initial conditions. In the early/onset stage, we assume that a small random number of normal cells become cancerous by mutation. Thus, initially, we assume that all biochemicals of notch signaling in each cell are at a resting state with a small perturbation (Fig. 2A). Namely,

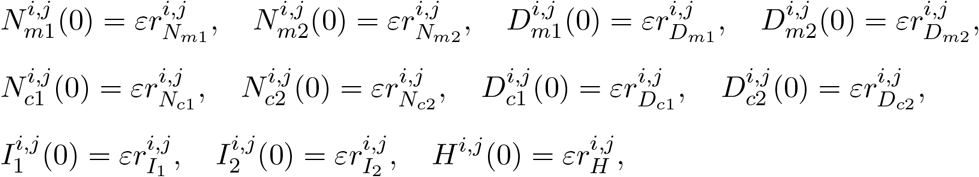

where *ε* = 0.01, 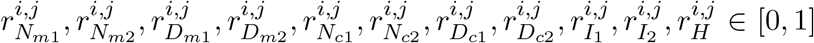 for 1 ≤ *i* ≤ *m* and 1 ≤ *j* ≤ *n*. On the other hand, for late stage, we assume a scenario of cancer stage 3 so that a mass of cancer of a certain size already exists. The detailed setting will be given in Section 3.5.

We also assume zero boundary conditions for all *t* ≥ 0 such that

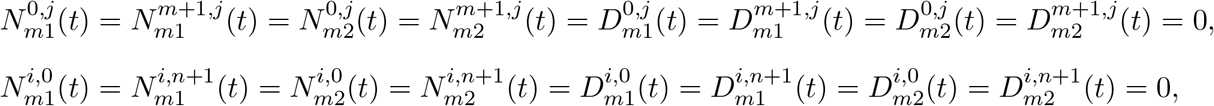

for 1 ≤ *i* ≤ *m* and 1 ≤ *j* ≤ *n*.

Sensitivity analysis is used to investigate how much model parameters affect the dynamics of the network. We employed the Extended Fourier Amplitude Sensitivity Test (eFAST) for this purpose (Saltelli et al. 1999; Seirin-Lee et al. 2020). In brief, we define a sensitivity function whose dynamics reveal the amount of HES-1 expression and track its changes in response to changes in parameter values. The sensitivity function is denoted by *F*_*S*_ and defined as

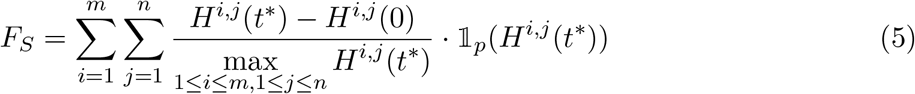

where

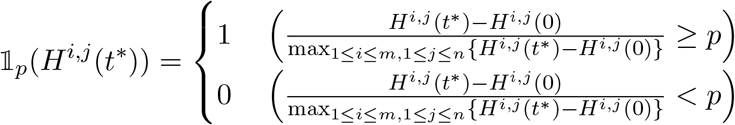

where *t*^*^ is the time scale at which proteins or the HES-1 gene have approached steady states. This sensitivity function provides a quantitative measure of how many cells exhibit increased expression of HES-1 in response to changes in parameter values.

## 3 Results

### 3.1 Intracellular signaling pathways that enhance a HES-1 expression in a single cell case

To explore which signaling pathways are most crucial in promoting cellular oncogenesis at a single-cell level, we first examined HES-1 expression levels in two cell cases with changing network intensities within certain ranges (Fig. 2B). We initially observed that variations in HES-1 expression are not sensitive to parameters related to the binding rates of Notch and Delta in the cell membrane (*α*_1_, *α*_2_, *α*_3_, and *α*_4_), the translocation rate of cytosolic Notch/Delta to the membrane (*γ*_*N*_ */γ*_*D*_), Delta production rates (*β*_41_ and *β*_42_), and the inhibition intensity of HES-1 expression by NICD (*ν*_3_), although certain parameter ranges exhibited a step-like variation in HES-1 expression (Fig. 3A). In contrast, Notch1 activation by NICD1 (*β*_21_) and HES-1 activation by NICD1 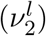 notably affected the variations of HES-1 in NSCLS type. Similarly, Notch2 activation by NICD2 (*β*_22_) and HES-1 activation by NICD2 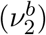 affected notably the variations of HES-1 in EBT type (Fig. 3A). To confirm these observations, we also carried out eFAST sensitivity analysis using the sensitivity function (Eq. (5) (Fig. 3B) and the results showed consistent outcomes. Taken together, these results indicate that a straightforward signaling pathway to increase HES-1 is most critical at the single-cell level in both NSCLS type and EBT type, and that the contrasting network pathways in NSCLS type and EBT type may be a crucial factor preventing simultaneous treatment. Furthermore, we could not find any other notable signaling pathways by which we can regulate HES-1 expression simultaneously in a single-cell case.

**Figure 3:**
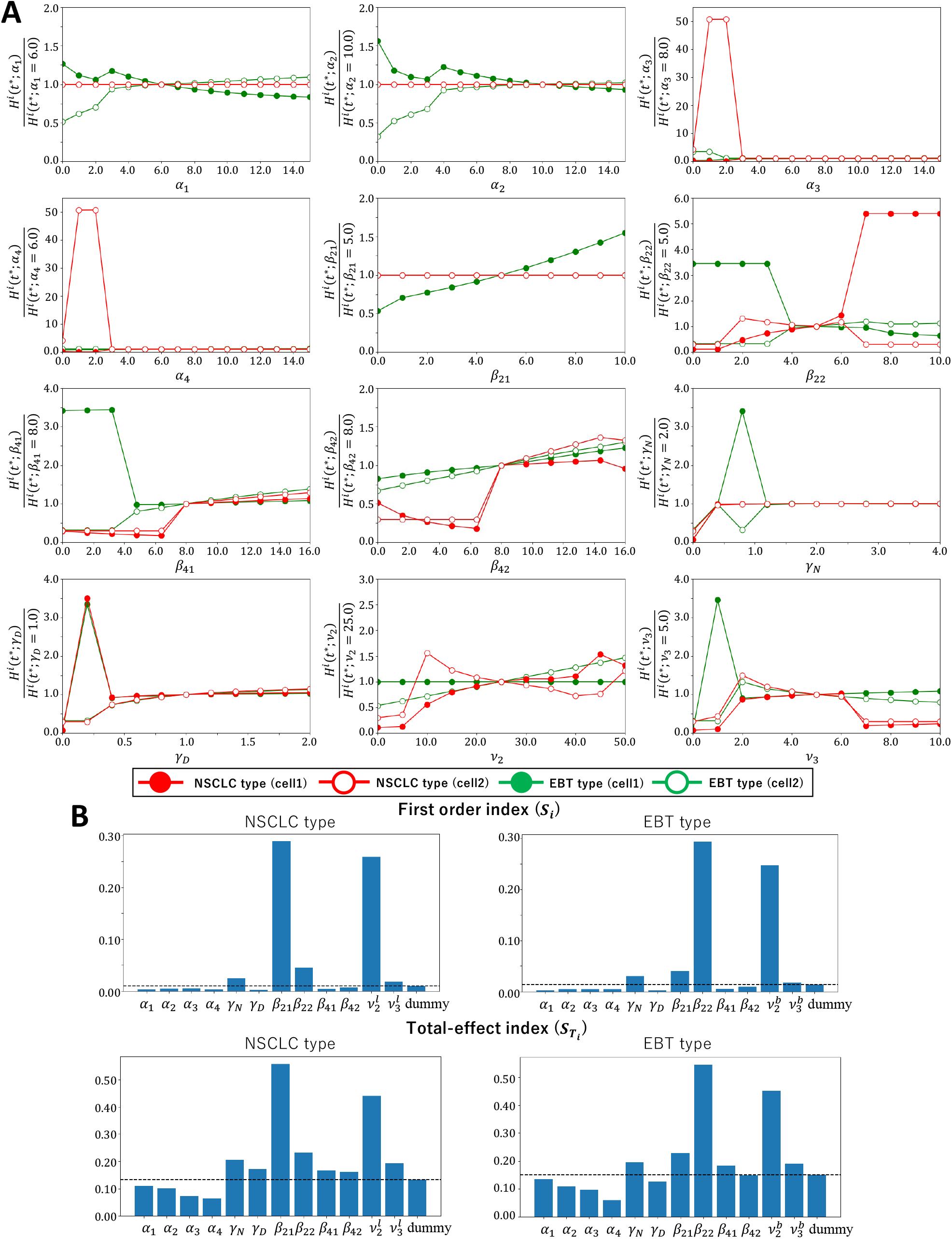
Effect of networks for a single cell case. (A) The increase rate of HES-1 as changing parameter values in each cell. The green line shows the results for the NSCLC type, and the red line shows the results for the EBT type. (B) The results of the sensitivity analysis. The dotted line represents the values of the dummy parameter.

### 3.2 Signaling pathways enhancing cancer cells with high HES-1 expression in multicellular system

In the previous analysis at the single-cell level, we confirmed the critical role of the contrasting signal pathway structure of Notch signaling in up-regulating HES-1 in each cancer type.

However, we were not able to identify other suitable signal pathways capable of simultaneously reducing HES-1 expression. Therefore, we next investigated whether it is possible to find a common signal pathway that reduces the total cancer cell number with high HES-1 expression in a multicellular system.

As shown in Fig. 4A, we found that Notch1-Delta1 binding (*α*_1_), Notch1-Delta4 binding (*α*_2_), Notch1 activation (*β*_21_), Notch2-Delta1 binding (*α*_3_), Notch2-Delta4 binding (*α*_4_), and Notch1/2 activation rate (*β*_21_/*β*_22_) led to contrasting tendencies in cell increase/decrease depending on the cancer types. In contrast, the translocation rate of Notch (*γ*_*N*_), translocation rate of Delta (*γ*_*N*_), and HES-1 activation/inhibition rate (*ν*_2_*/ν*_3_) showed the similar tendency of cancer cell number variations between the two cancer types. In particular, the cases of translocation rates of Notch/Delta showed a decreasing tendency in cancer cell numbers as the network intensity increased, indicating that these two networks could be targeted to suppress the increase of cancer cells in both cancer types that have contrasting Notch networks.

**Figure 4:**
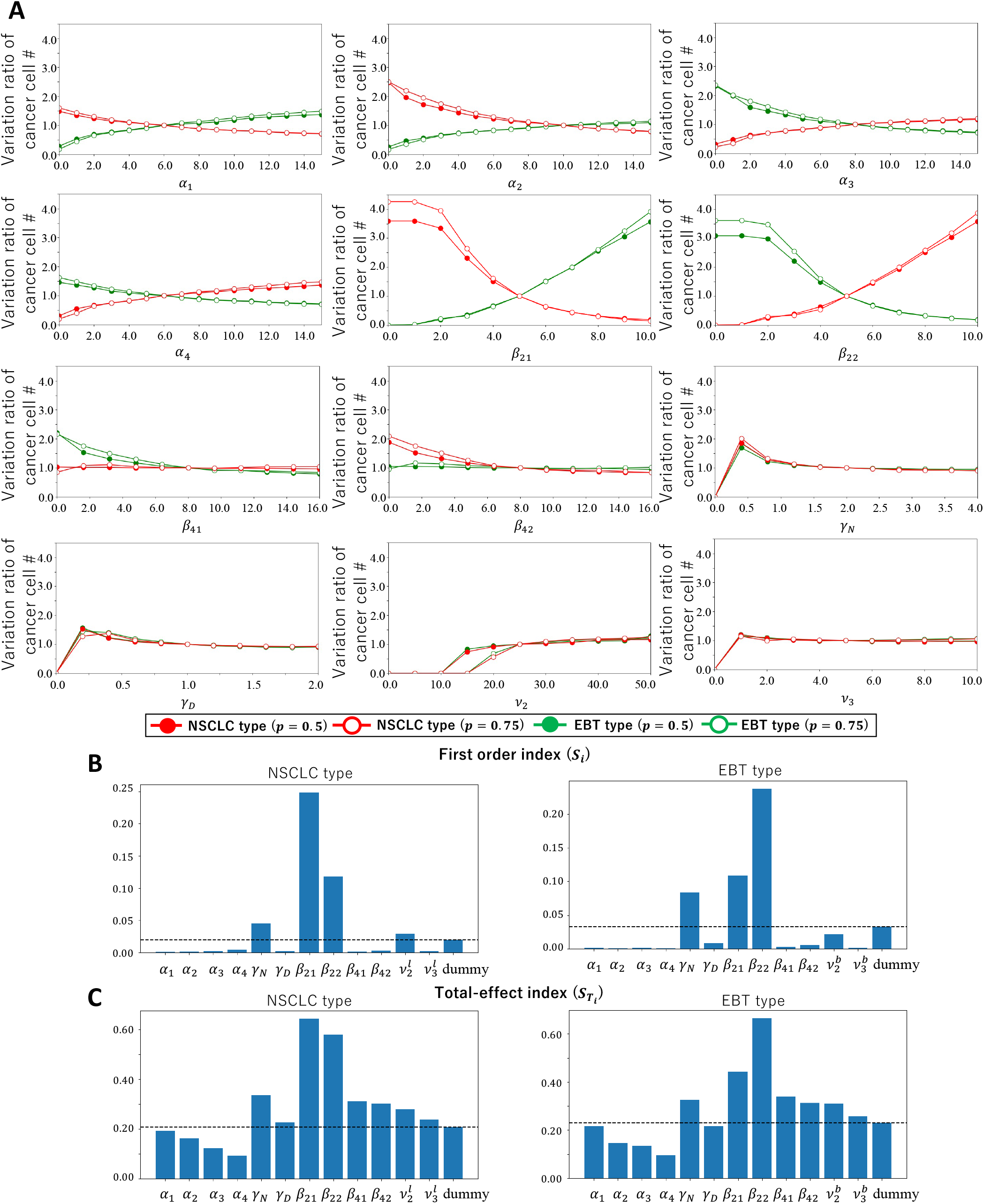
Effect of networks for multicellular system case. (A) The variation ratio of cancer cell number. Eq. (4) was calculated. (B-C) The results of sensitivity analysis based on the sensitivity function (5) with 100 × 100 cells The dot line represents the first order index or total-effect index of dummy parameter.

To confirm the simulation results of representative parameter sets, we conducted sensitivity analysis using the sensitivity function (5) (Fig. 4B) with 3746 parameter sets. Excluding the direct activation networks of HES1 (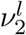 and 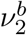), we found that three parameters, the Notch1/2 activation rate (*β*_21_/*β*_22_) and the translocation rate of Notch (*γ*_*N*_), showed high indices in both first-order and total effect indexes for both cancer types. We concluded that the decrease of both Notch1 and Notch2 (*β*_21_ and *β*_22_) or the translocation rate of Notch (*γ*_*N*_) is the most suitable targeted network for reducing cancer cells in both cancer types.

### 3.3 Network cleavage/enhancement therapy tests

Based on the network analysis in the previous section, we next tested the effectiveness of therapies. We first tested network cleavage therapies for monotherapy and combination therapies by using the model (2); *γ*_*N*_ -cleavage (S1 cleavage), Notch-Delta binding-cleavage (S2/3 cleavage), our new suggested *β*_21_*/β*_22_-cleavage (S^*new*^ cleavage) (Fig. 2C). S1 cleavage suppresses the network of translocation of notch which we found as a targeted network for both cancers. S2/3 cleavage is directly related with cutting inter-cellular network. S^*new*^ cleavage reduces the activations of both Notch1 and Notch2 by NICD1 and NICD2 at the same time. As an evaluation measure, we counted relative cell numbers expressing high HES-1 (𝒩 (*C*_*p*_)_*treated*_*/𝒩* (*C*_*p*_)_*untreated*_) as defined in Eq. (4). We also calculated it with 10 random samples of therapy intensities for cleavage and enhancement therapies, respectively, namely, 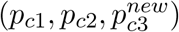 and (*p*_*e*1_, *p*_*e*2_, *p*_*e*3_), to consider the case where variations in therapy intensities exist between patients.

First of all, we found that monotherapy is almost ineffective for both cancer types, although S1 cleavage therapy did not result in an increase in cancer cells. In particular, some therapies even increased the number of cancer cells, as shown in Fig. 5A. Interestingly, however, the combination of the three therapies decreased the number of cancer cells by approximately 20% and showed notable effectiveness.

**Figure 5:**
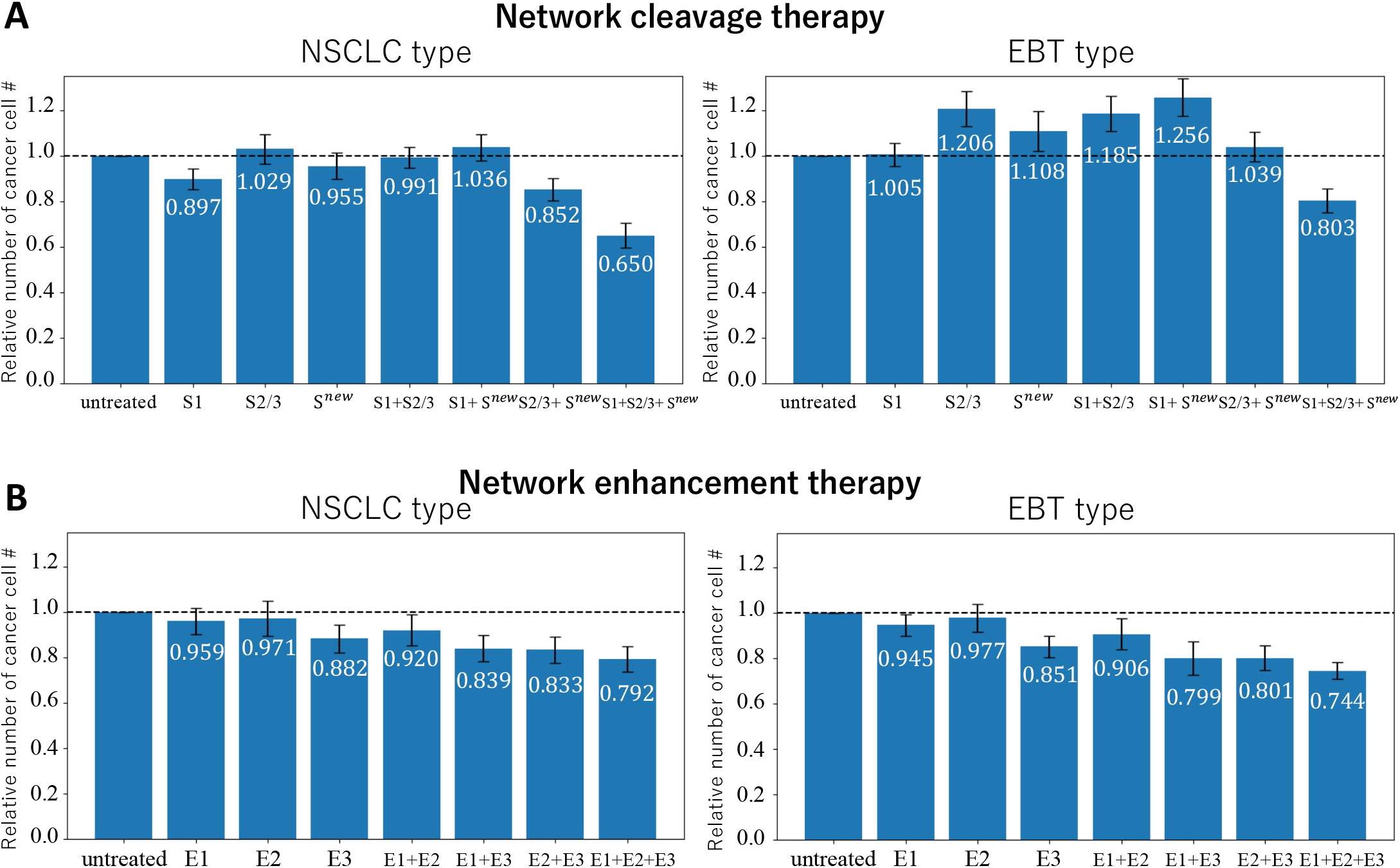
*in silico* therapy tests. (A) The results of network cleavage therapy for ten random sets of (*p*_*e*1_, *p*_*e*2_, and *p*_*e*3_). (B) The results of network enhancement therapy for ten random sets of (*p*_*e*1_, *p*_*e*2_, and *p*_*e*3_). The graphs are plotted by average with standard deviation. The exact values of average are shown in the bar graph. The simulation tests have been done for 100 × 100 cells.

Since most of the mono/combination cleavage therapies even increased the number of cancer cells, we next tested the network in the opposite way, namely, enhanced therapies with the model (3); E1 enhancement, E2 enhancement, and E3 enhancement (Fig. 5B). Interestingly, we found that the enhanced therapies are more effective than cleavage therapies, especially concerning mono-therapies and two-types combined therapies. However, the maximal effectiveness of the three-types combined therapy was not notably different between cleavage and enhanced cases. Taken together, both cleavage and enhanced therapies can reduce the number of cancer cells most effectively with the three-types combined therapy.

### 3.4 Effectiveness of therapy order

We next asked whether the order of therapy combinations is important in increasing the effectiveness of therapies. We tested two cases: one where treatment is administered every day (Continuous treatment) and the other where treatment is administered only once with different therapy (Discontinuous treatment). We determined the timing to start the next treatment based on when the cancer cell numbers approached a steady state approximately, every 10 months ∼ one year (Lamarca et al. 2019). As shown in Fig. 6, we initially observed that discontinuous treatment is less effective than continuous treatment. Furthermore, there were instances where an increase in cancer cells occurred (Fig. 6A, Discontinuous treatment, green lines, red line marked white circle, and blue line marked blue circle). Interestingly, the network enhancement therapies were no longer effective compared to the network cleavage therapies (Fig. 6A, B, Continuous treatment), indicating that the combination order can critically affect treatment effectiveness.

**Figure 6:**
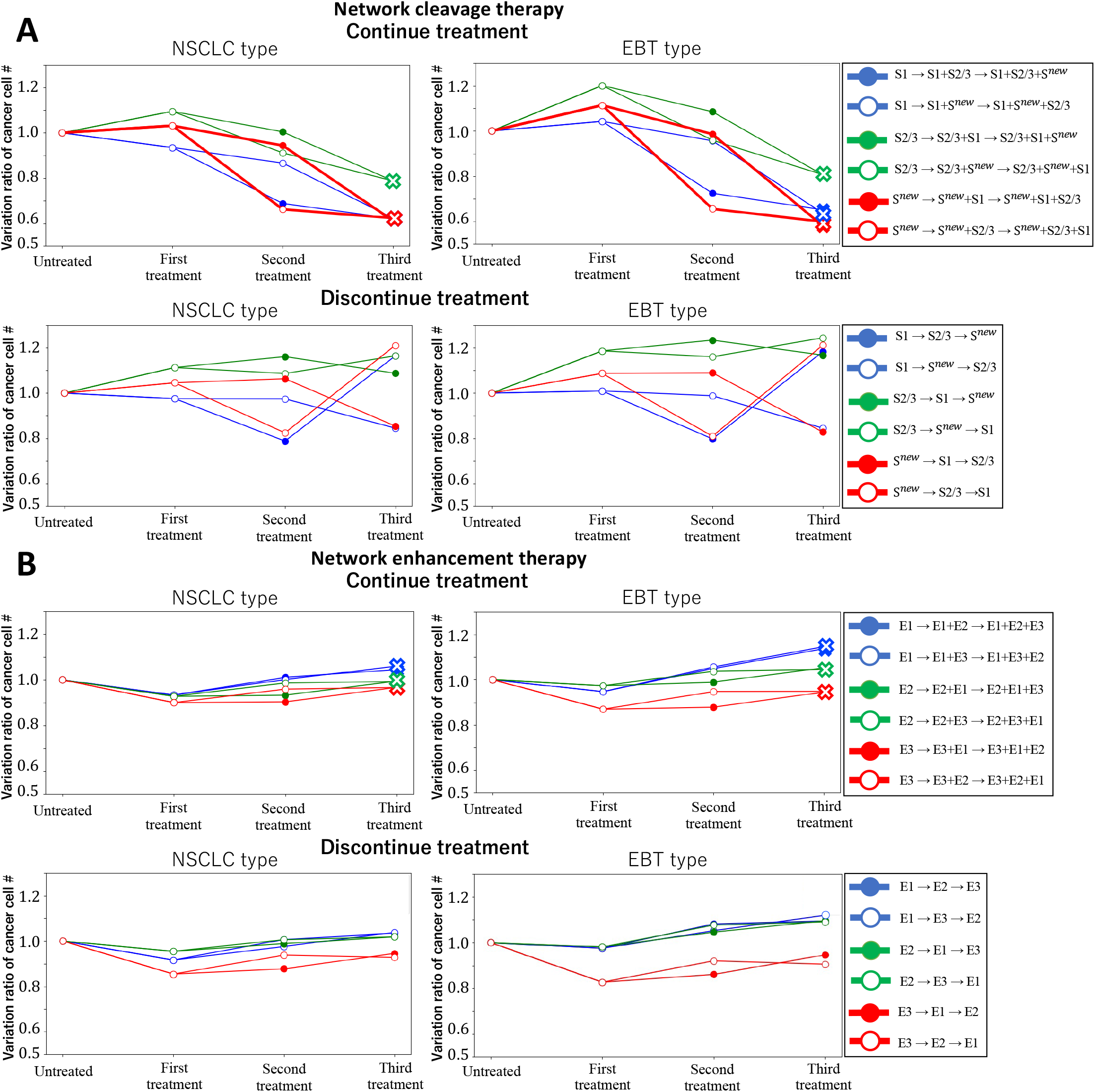
The effect of the order of treatment on cancer cell number. (A) The results of network cleavage therapy. (B) The results of network enhancement therapy. The number of cancer cells was counted after each therapy was administered and divided by the number of cells in the untreated case. The simulation tests have been done for 100 × 100 cells.

Next, we precisely examined which combination orders are most effective in the continuous treatment of network cleavage therapies. We found that the combination starting with S^*new*^ therapy exhibits the highest effectiveness after the third treatment (Fig. 6A, continue treatment, red lines). Moreover, the ratio of cancer cells after all treatments was much lower than in the case of simultaneous treatments (Fig. 6A, continue treatment, red line marked x symbol). Additionally, treating S2/3 therapy followed by S1 therapy showed a more effective decrease in cancer cells than the opposite order. That means that selecting a therapy order that achieves significant effectiveness early on is feasible, potentially reducing the mental burden of treatments for patients. Taken together, we concluded that the combination order of S^*new*^ → S2/3 → S1 could be the most optimal therapy for both NSCLS type and EBT type.

### 3.5 Optimal index for effective treatments on cancer spreading

In the previous sections, we explored the effectiveness of therapies under a scenario where cancer cells develop simultaneously in a local region, neglecting a spreading case from a given size of cancer cell mass. Here, we assume a scenario where an 3 stage of cancer is detected initially, and we evaluate not only the number of cancer cells but also the spreading speed on therapy effectiveness.

To this end, we set up a cancer cell mass with a high level of HES-1 expression in the center of the tissue and gave a sufficiently low level of HES-1 cells surrounding it, based on the actual size of single cells, single cancer cells, and the sizes of the lung and cerebellum (Harouaka et al. 2013)(Fig. 7A). We then started a simulation with permitting Delta-Notch binding when one of the neighboring cells expresses higher HES-1 than a given standard level as follows (Fig. 7B):

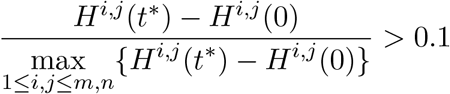

**Figure 7:**
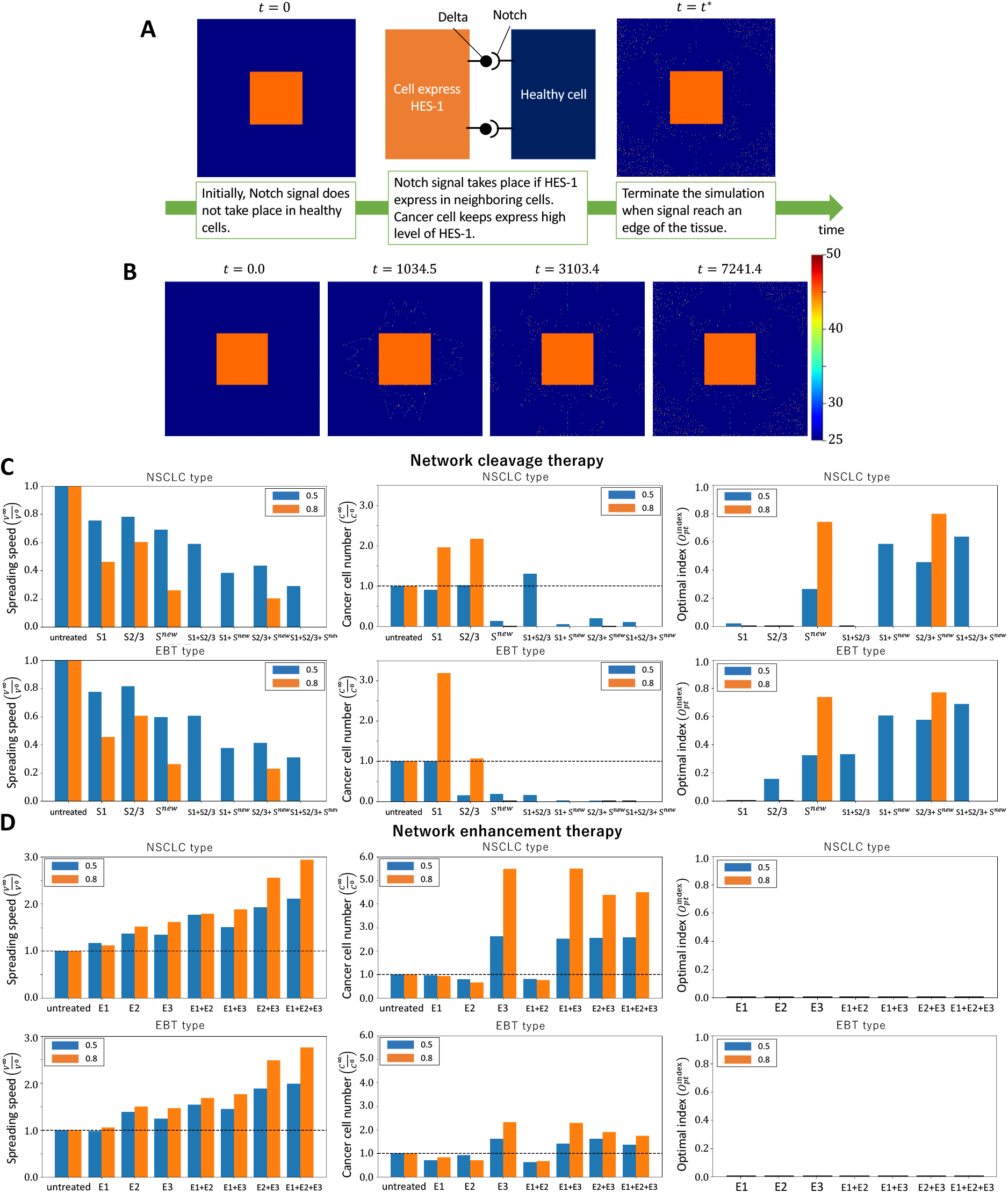
*in silico* test of treatments on cancer spreading. (A) The method of simulation on cancer spreading. (B) An example of simulation results on cancer spreading without treatment in NSCLC type. (C) The results of network cleavage therapy. Blue bar shows the results for 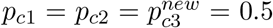. Orange bar shows the results for 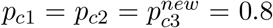. (D) The results of network enhancement therapy. Blue bar shows the results for *p*_*e*1_ = *p*_*e*2_ = *p*_*e*3_ = 0.5. Orange bar shows the results for *p*_*e*1_ = *p*_*e*2_ = *p*_*e*3_ = 0.8. The number of cells has been set as 1000 × 1000, with the initial number of cancer cells being 332 × 332 of those.

(The more details of simulation conditions are given in Appendix A). We next defined the optimal index of treatment by

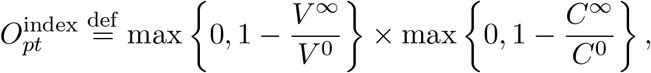

where

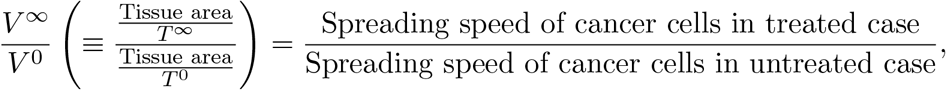

*T* ^∞^ and *T* ^0^ are the time for the treated/untreated case when cancer cells approached the boundary of tissue.

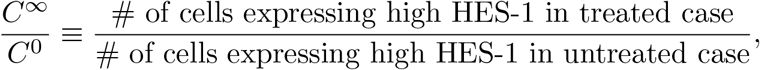

where we counted cell numbers except for the initial cancer cell mass. The measures above indicate that if *V* ^∞^*/V* ^0^ *<* 1 and *C*^∞^*/C*^0^ *<* 1, the therapy is considered effective. Otherwise, the therapy is deemed ineffective.

For the network cleavage therapy, we first found that all therapies are effective in decreasing the spreading speed of cancer cells but are not all effective in terms of reducing cell numbers, as shown in Fig. 7C left panels and center panels. On the other hand, for the enhancement therapy, most therapies showed very ineffective results, also shown in Fig. 7D left panels and center panels, indicating that cleavage therapy should be chosen for effective treatments.

Finally, we calculated the optimal index of treatments (Fig. 7C, D, right panels) and found that both NSCLS type and EBT type can be treated effectively with the same targeted therapies such as S^*new*^, S1+S^*new*^, S2/3+S^*new*^, and S1+S2/3+S^*new*^. Furthermore, both the monotherapy case and combination therapies with our suggested new therapy showed a higher optimal index of treatments. This suggests that the network cleavage of Notch1/2 activation by NICD1/2 in cytosol could serve as a core targeted therapy capable of yielding optimal treatment outcomes in both cancers, despite their contrasting notch signaling pathways.

## 4 Discussion

In this study, we explored whether there exists an effective targeted therapy to suppress the expression of the oncogene HES-1 in two types of cancers based on contrasting notch signaling pathways. In single-cell investigations, where two cell interactions of notch signaling were set, a direct activating network to HES-1 was identified as the most critical up-regulation concerning each contrasting notch signaling pathway. Thus, we could not find any clues for a targeted therapeutic network through single-cell analysis. However, multi-cellular *in silico* experiments revealed that the translocation network of notch in each cell could serve as a targeted network to reduce the expression of HES-1, even in both contrasting structures of notch signaling. Furthermore, this finding suggests that a key network suppressing HES-1 exists not only in inter-cellular networks but also in intra-cellular networks, even within inter-cellular interacting-based notch signaling pathways.

Based on the results of sensitivity analysis on network intensities related with HES-1 up-regulation, we next have explored the effectiveness of therapeutic tests with mono/combinations cases of existed target therapies and our new suggested therapies. In general, mono-therapy was not effective to decrease the cancer cells and multiple combination of therapy showed a better effectiveness. We also compared the effectiveness for the cases of simultaneous and sequential treatments. In the case of simultaneous treatments, the full combination of the three therapies is the most effective in both cleavage and enhanced therapies.

On the other hand, the sequential treatments showed completely different effectiveness between cleavage and enhanced therapies. Even though any order combinations, the enhanced therapy could not be an optimal treatment to reduce cancer cells effectively. On the other hand, the sequential treatment starting from the cleavage therapy of the network from Notch to NICD (S^*new*^ therapy), which is newly suggested in this study, showed more effective reduction of cancer cells than simultaneous combination treatment case of the three therapies and S2/3 therapy followed by S1 therapy was the most effective order.

Finally, we also evaluated the effectiveness of therapies with both the decrease ratio of cancer cell number and spreading speed of cancer cells. We again found that the enhanced therapy is not effective and the cleavage therapies of S^*new*^ therapy and its combination with S2/3 therapy are most effective on both cancers. This result suggests that a signal blocking in not only intra-cellular network but also inter-cellular network is crucial to suppress the expansion of cancer cells. In fact, the network of S2/3 therapy was not detected as an important network to up-regulate HES-1 in our sensitivity analysis. This implies that the spatial conditions/initial situation of cancer before treatment should be considerable factor to choose an optimal therapy. In this study, we have focused on suggesting an optimal targeted therapy to suppress cancer cells of which mechanisms are based on the contrastive notch signaling pathway. Our results showed that multiple cancers based on two contrasting Notch networks could be controlled by a common targeted signal network. A combination therapy with multiple targeted agents enhancing Notch production in the cytosol, binding of Notch-Delta in the membrane, and transporting Notch to the membrane may be most effective in the early stages of cancer development. In contrast, in the late stages, cleavage therapies may be more effective in suppressing cancer expansion. We further found that the therapy order of treatments is very critical on the effectiveness of reduction of cancer cells. All these results suggest that optimal treatment should be sensitive to the stage of cancer development, and the choice of targeted therapy should be carefully considered depending on the cancer stage with the optimal combination and order of treatments. On the other hand, we did not consider the scenario of Notch signaling-induced cell division in this study (Ranganathan et al. 2011), although the high proliferation rate of cancer cells is directly expected to affect the increase in cancer cell numbers proportionally with the number of high levels of HES-1 cells. Recently, we developed a mathematical model of the Notch signaling system, including the effect of cell division, and this study is in progress.

## Data availability

All relevant data are included in the manuscript and the supporting information files.

## Code availability

Numerical code is made available online on GitHub (https://github.com/TWakamoto/Notch-signaling.git) and all other data are available from the authors (TW and SSL) on reasonable request.

## Author Contributions

SSL initiated and designed the research. SSL and TW developed a mathematical model and executed the research. TW developed numerical algorithms and executed simulations. SSL and TW analyzed data. TW wrote the first draft, and SSL significantly revised the manuscript.

## Acknowledgements

The author would like to express special thanks to Dr. Hiroshi Ishii (Hokkaido University) and Mr. Michito Ujino for their valuable discussion and comments. In particular, we would like to thank to Dr. Hiroshi Ishii and Dr. Steffen Plunder for giving a great assistant for improving programming codes. This study was supported by the Japan Science and Technology Agency(JST) CREST (JPMJCR2111) and MEXT, Grant-in-Aid for Transformative Research Areas(A) (22H05110) to SSL.

## A Appendix Details of simulation conditions for cancer cell spread case

Initially, healthy cells are steady state and Notch signal does not take place. Notch signal starts if one of neighboring cells express more than standard level of HES-1 as follows:

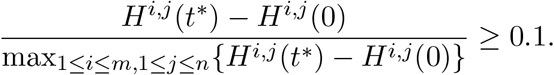

Cancer cells keeps to express high level of HES-1. Finally, the simulation is terminated when signal reach one of the boundary of the tissue.

Let Ω ⊂ ℝ ^2^ be the region in which Notch-Delta signaling takes place. We also define the region consisted by cells with high level of HES-1 and the region consisted by cells expressing low level of HES-1 at *t* = 0 by Ω_*h*_ ⊂ Ω, and the region consisted by cells expressing low level of HES-1 at *t* = 0 by Ω_*l*_ ⊂ Ω which is complement of Ω_*h*_ i.e. Ω_*l*_ := Ω *\* Ω_*h*_. The dynamics of the biochemical in the cell (*i, j*) ∈ Ω_*l*_ expressing low level of HES-1 at first can be written as follows:

### Notch in membrane

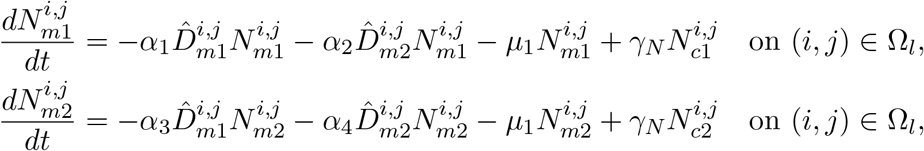

### Delta in membrane

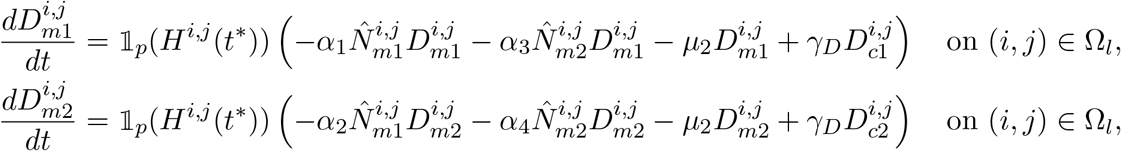

### NICD in cytosol

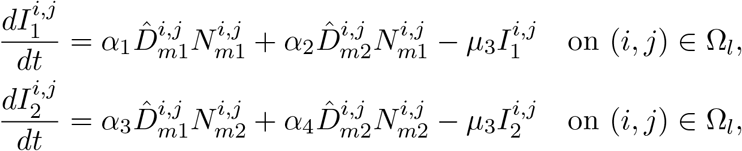

### HES-1 in nucleus

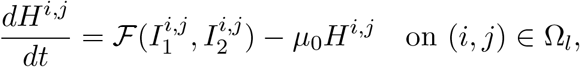

### Notch in cytosol

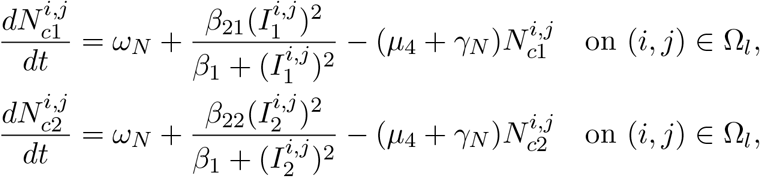

### Delta in cytosol

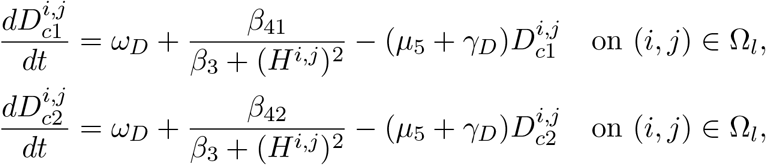

where

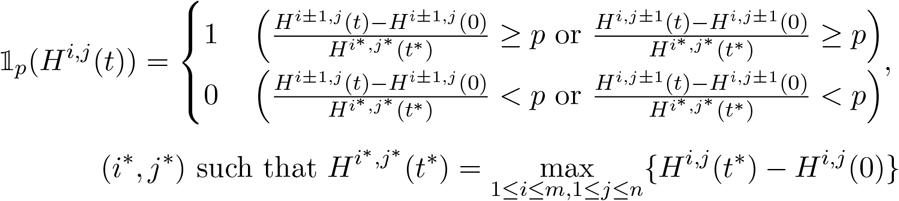

We did not considered basal production of Notch and Delta in other simulation in this paper since basal production is very small. However, we consider basal production in the simulation of cancer spreading. *ω*_*N*_ and *ω*_*D*_ are basal production of Notch and Delta in cytosol, respectively.

Since the cell (*i, j*) ∈ Ω_*h*_ keeps to express high level of HES-1 constantly, the dynamics can be written as follows:

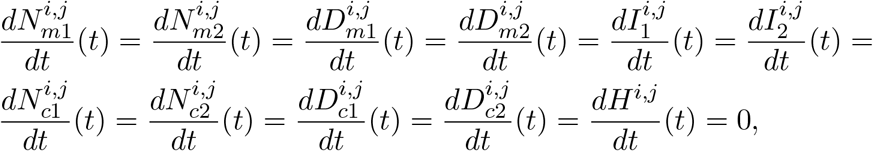

for any (*i, j*) ∈ Ω_*h*_ and *t >* 0.

The initial condition has been given as

### Notch in membrane

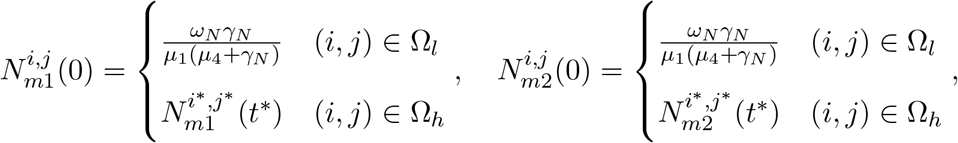

### Delta in membrane

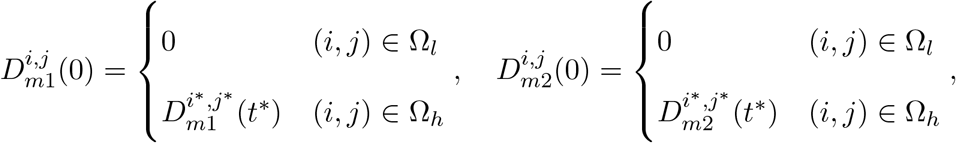

### NICD in cytosol

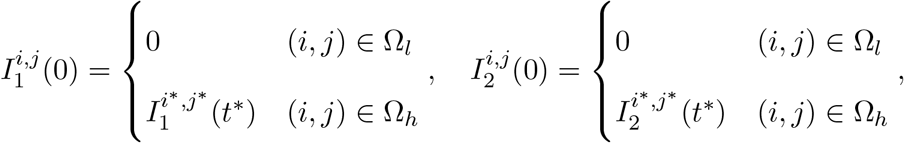

### HES-1 in nucleus

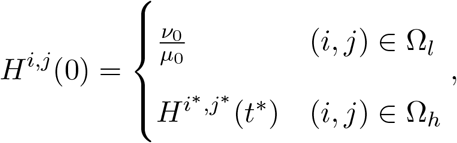

### Notch in cytosol

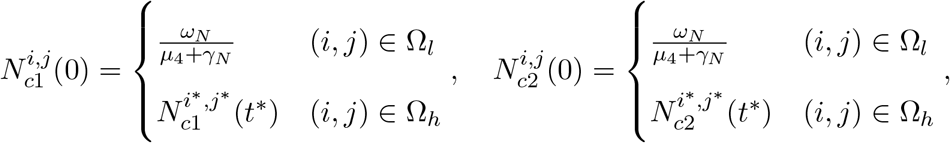

### Delta in cytosol

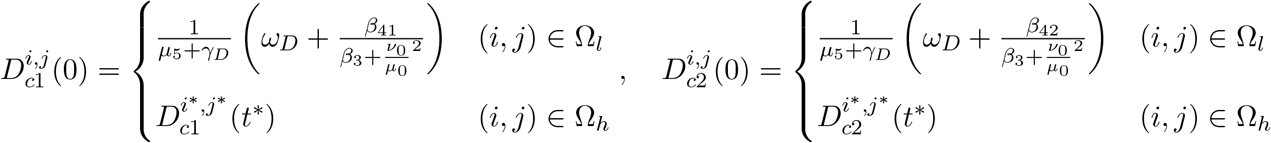

In the area Ω_*l*_, we set the initial condition such that the dynamics of biochemicals is steady state automatically. The boundary condition has been given same as the simulation in section 3.2.

